# Localized structural frustration for evaluating the impact of sequence variants

**DOI:** 10.1101/052027

**Authors:** Sushant Kumar, Declan Clarke, Mark Gerstein

**Affiliations:** Program in Computational Biology and Bioinformatics, Yale University; Department of Molecular Biophysics and Biochemistry, Yale University; Department of Chemistry, Yale University; Department of Computer Science, Yale University, 260/266 Whitney Avenue PO Box 208114, New Haven, CT 06520, USA

## Abstract

The rapidly declining costs of sequencing human genomes and exomes are providing deeper insights into genomic variation than previously possible. Growing sequence datasets are uncovering large numbers of rare single-nucleotide variants (SNVs) in coding regions, many of which may even be unique to single individuals. The rarity of such variants makes it difficult to use conventional variant-phenotype associations as a means of predicting their potential impacts. As such, protein structures may help to provide the needed means for inferring otherwise difficult-to-discern rare SNV-phenotype associations. Previous efforts have sought to quantify the effects of SNVs on structures by evaluating their impacts on global stability. However, local perturbations can severely impact functionality (such as catalysis, allosteric regulation, interactions and specificity) without strongly disrupting global stability. Here, we describe a workflow in which localized frustration (which quantifies unfavorable residue-residue interactions) is employed as a metric to investigate such effects. We apply frustration to study the impacts of a large number of SNVs available throughout a number of next-generation sequencing datasets. Most of our observations are intuitively consistent: we observe that disease-associated SNVs have a strong proclivity to induce strong changes in localized frustration, and rare variants tend to disrupt local interactions to a larger extent than do common variants. Furthermore, we observe that somatic SNVs associated with oncogenes induce stronger perturbations at the surface, whereas those associated with tumor suppressor genes (TSGs) induce stronger perturbations in the interior. These findings are consistent with the notion that gain-of-function (for oncogenes) and loss-of-function events (for TSGs) may act through changes in regulatory interactions and basic functionality, respectively

## Introduction

The advent of next-generation sequencing technologies has led to a remarkable increase in genomic variation data at both the exome as well as the whole-genome levels ^1,2^. These large datasets are playing a pivotal role in advancing efforts toward personalized medicine ^3^. Non-synonymous coding single nucleotide variants (termed SNVs throughout this study) are of particular interest because of their implications in the context of human health and disease ^4–6^. As such, considerable effort has been invested in curating disease-associated SNVs into various databases, including the Human Gene Mutation Database (HGMD) ^5^, ClinVar ^6^and the Online Database of Mendelian Inheritance in Man (OMIM) ^4^. Concurrently, initiatives such as The 1000 Genomes Project ^7,8^, Exome Sequencing Project (ESP) ^9^ and Exome Aggregation Consortium (ExAC) ^10^ have generated large catalogues of SNVs within individuals of diverse phenotypes.

As the costs associated with sequencing entire human genomes and exomes continue to fall, sequencing will become routine in both medical and academic settings ^11^. Indeed, it may take less than a decade to reach the milestone of a million sequenced genomes ^12^, resulting in massive datasets of rare SNVs. This exponential growth in the number of newly discovered rare SNVs poses significant challenges in terms of variant interpretation ^13^. Compounding this challenge is the fact that many of these variants will be unique to single individuals. The extremely low allele frequencies of such “hyper-rare” SNVs render them too rare to draw variant-phenotype associations with confidence – unlike more common variants, the very rarity of these ultra-rare genomic signatures renders phenotypic inference through association studies extremely difficult. Together, these trends underscore a growing and urgent need to evaluate the potential effects of low-allele-frequency variants in unbiased ways using high-throughput methodologies.

Simultaneously, immense progress has been made in resolving the three-dimensional structure of many proteins over the last several decades ^14^. A large volume of high-resolution data on protein-protein, protein-ligand and protein-nucleic acid complexes is now available. This complementary evolution of sequence and structural databases provides an ideal platform to investigate the functional and structural consequences of benign and disease-associated SNVs on protein structures. The integration of variant and structure knowledge bases will lead to a greater understanding of the biophysical mechanisms behind various diseases. In addition to gaining a better understanding of how disease-associated SNVs impart deleterious effects, this integration can be utilized to both predict the impacts of poorly understood SNVs (i.e., SNVs which are known to be deleterious, but for which a plausible biophysical or functional rationale is missing) and to prioritize SNVs based on predicted deleteriousness ^15–18^. We also note that this approach may aid in more intelligent and targeted design of drugs in various therapeutic contexts.

In last few decades, many studies have evaluated the impacts of SNVs by examining or predicting changes in thermodynamic stability ^19–21^. These approaches rely on the fact that SNVs may induce substantial changes in the folding landscape and conformational ensemble. Such changes in global stability are often quantified by calculating the folding free energy change (ΔΔG) after mutating residues ^21,22^.Importantly, however, many disease-associated SNVs introduce local structural changes without appreciably affecting folding free energy or global stability ^23,24^. Such local perturbations may include disruptions in residue packing or hydrogen bond networks ^25,26^ and salt bridges ^27,28^. Examples of the associated effects include disruptions to catalytic centers, changes to “hotspot residues” that are responsible for interaction affinities and specificity, as well as perturbations to key allosteric sites ^29–31^. Changes to such residues may impart only minimal effects to the protein’s overall topology, but may nevertheless drastically influence protein behavior and functionality.

We examine the role of localized perturbations by calculating changes in the localized frustration indices(termed frustration throughout this study) ^32,33^ of residues impacted by SNVs. Qualitatively, the frustration of a given residue quantifies the degree to which the residue is involved in favorable or unfavorable interactions with neighboring residues in space. The residue change that is introduced by an SNV may result in more (less) unfavorable interactions with neighboring residues, thereby increasing (decreasing) the frustration at that site. SNVs thereby act as agents that may relieve unfavorable interactions or alternatively impair local stability, depending on the nature of the amino acid substitution and the surrounding environment within the protein. Throughout this study, such changes in frustration are designated by ΔF.

The concept of frustration was originally introduced by Wolynes *et al.* to describe the protein folding landscape ^32^. The protein folding process is believed to follow a smooth funneled energy landscape, in which strong energetic conflicts are avoided ^34–38^. However, despite minimizing configurations that exhibit frustration, local frustration is essential to protein biology and function ^39–41^. Highly-frustrated local interactions result in micro-states of high potential energy. Such micro-states provide proteins with the avenues needed to carry out essential functions that entail a release of energy and the concomitant shifts in occupied energetic wells. Examples of processes that require these “energetic bursts” include catalysis, allosteric communication, conformational switches and proteinquakes ^42^, as well as protein-protein interactions ^32,43,44^. Ferriero et al. proposed a framework to compute the frustration profile of a given protein (32). The localized frustration index quantifies the contribution of each residue or residue pair in the total energy of the native structure compared to their energetic contribution in a random non-native configuration (see Methods and (45)). A native residue (residue pair) is considered to be minimally frustrated if it contains sufficient extra stabilization energy in its native state. In contrast, a sufficiently destabilizing residue (residue pair) in the protein structure is considered to be maximally frustrated ^45^. In addition, a residue (residue pair) is considered to be neutral when its stability profile lies between these extremes.

We take a data-drive approach to analyze ΔF profiles produced by the introduction of SNVs in a large dataset of proteins. SNVs present in healthy human populations (The 1000 Genome and ExAC projects) are highly enriched in benign SNVs. Therefore, we term SNVs in these datasets as “benign” (though we qualify this term by noting that a small subset of these SNVs may actually impart as yet undetected deleterious effects). However, within these datasets, there are various degrees along the continuum of phenotypic effects. While deleterious variants are more enriched among rare SNVs, neutral variants have stronger representation among common variants. In addition, we also quantified and compared ΔF profiles introduced by disease-associated SNVs (these SNVs were taken from the HGMD database), as well as cancer somatic variants, thereby enabling in-depth analyses of the differential effects between SNVs in driver and passenger genes.

The majority of our analyses were consistent with prior studies investigating how SNVs impact protein structures, we provide a distinct rationale through the lens of localized frustration. We observe that large disruptions in local interactions of minimally frustrated core residues distinguishes disease-associated SNVs from benign SNVs as well as SNVs impacting driver and passenger genes in cancer. In contrast, benign SNVs in passenger genes generate larger perturbations in local interactions of minimally frustrated surface residues compared to core residues. Furthermore, comparisons between rare and common SNVs within healthy human populations indicate that rare variants induce larger disruptions in favorable local interactions compared to common variants. Moreover, we also investigated the effects of SNVs impacting conserved and variable regions of proteins, where conservation was measured across different species. For disease-associated SNVs, we detected a significant disparity between local perturbations observed due to SNVs impacting conserved regions compared to variable regions of proteins. However, no such disparity was observed for benign SNVs.

We also demonstrate how frustration may provide insights in the context of oncogenes and tumor suppressor genes (TSGs). We find that somatic SNVs in oncogenes disrupt local interactions of surface residues and potentially facilitate cancer progression through the introduction of non-specific regulatory interactions. However, SNVs in TSGs drive cancer progression through larger local perturbations in core residues. These observations indicate that SNVs intersecting TSGs and oncogenes as having loss-of-function (LOF) and gain-of-function (GOF) effects, respectively.

## Results

### Differential effects of benign and disease-associated SNVs on ΔF profiles

We performed a comparative analysis to investigate the impacts of benign (1KG & ExAC) and disease-associated (HGMD) SNVs on the ΔF profiles of mutated residues in a large number of proteins. As detailed in Methods, each SNV dataset was divided into three distinct categories based on the frustration index of the wild-type residue. Maximally frustrated residues in the native structure exhibit conflicting interactions and unfavorable geometry in their local environment, thereby inducing local destabilization. Conversely, minimally frustrated residues are involved in biophysically favorable local interactions, and thus favorably contribute to the protein’s stability.

For each SNV, ΔF was calculated as follows (Figure 1). For a given SNV mapped to a PDB structure, two protein structures are used in our analysis: the native structure (as it exists in the PDB), and a model of the structure as it may exist when the affected residue is mutated (this is modeled by optimizing the structure after introducing the SNV). If a given SNV maps to residue location j within the structure, then within each of these two structures, the frustration index is calculated at residue j (the corresponding values are denoted as F_nat_ and F_mut_ for the native and mutated model structures, respectively). Subsequently, we determine the difference between the frustration index of the wild-type residue in the native structure and the mutated residue in the modeled structure (ΔF =F_mut_ – F_nat_).

**Figure 1:**
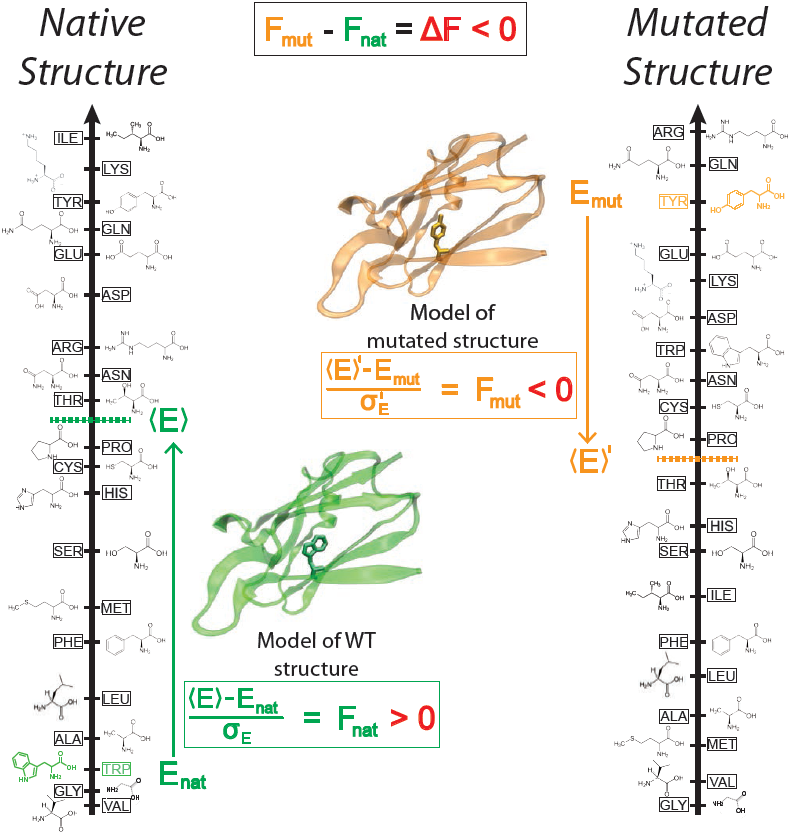
An example illustrating the case in which ΔF < 0. The ΔF associated with an SNV is negative if the SNV introduces a destabilizing effect. Shown here is the result of changing residue ID 31 in plastocyanin (pdb ID 3CVD) from the wild-type residue (Trp) to a mutated residue (Tyr). *Left)* The protein in its wild-type form (in green), in which the tryptophan residue at position 31 is substantially more energetically favorable relative to the mean energy ⟨E⟨) that would result from having any of the possible 20 amino acids at that position. This disparity is designated by (⟨E⟩ - E_nat_)/α_E_ = F_nat_ > 0. *Right)* The entire protein structure is then modeled (see methods) to generate the mutated structure after the SNV W31Y is introduced, thereby changing the relative energetic distributions for the different amino acids. The new mean and standard deviation associated with the energies of the modeled structure are designated by (E) and *OE*, respectively. In this case, the SNV that introduces 31Y results in an energy that is higher than the mean energy of all possible 20 amino acids at that position. This disparity is designated by (⟨E⟩’-E_mut_)/α_E_’ = F_mut_ < 0. Taken together, the negative value associated with the disparity between the F_mut_ and F_nat_ values (F_mut_ - F_nat_ = ΔF < 0) indicates that the this SNV is locally unfavorable.

After calculating the ΔF values in all three categories, the resultant distributions are plotted (further details are given under Methods). We observed that most SNVs (across all datasets) affecting maximally frustrated residues in the native structure induce small but positive ΔF values. This suggests that changes to maximally frustrated residues alleviate conflicting interactions, thereby resulting in a positive frustration difference (ΔF > 0). In contrast (and as expected), residues that are originally minimally frustrated tend to become more frustrated upon mutation, thereby, leading to a negative frustration difference (ΔF < 0) in majority of cases across each dataset. However, we emphasize that losses or gains in favorable interactions are dependent on the type of SNV (benign or disease-associated) as well as whether the SNV affecting a surface or core residue.

We observed that benign SNVs lead to greater disruptions within minimally frustrated surface residues compared to core residues in the native structure, and this trend is observed when using both ExAC and 1KG datasets (*p-value < 2e-16 from two-sample Wilcoxon test*) (Figure 2A & 2B). In addition, disease-associated SNVs (from HGMD) result in similar frustration changes between core and surface residues (*p-value < 2e-16 from two-sample Wilcoxon test*) (Figure 2C). However, SNVs from HGMD that impact minimally frustrated core residues induce stronger perturbations than benign SNVs influencing minimally frustrated core residues.

**Figure 2:**
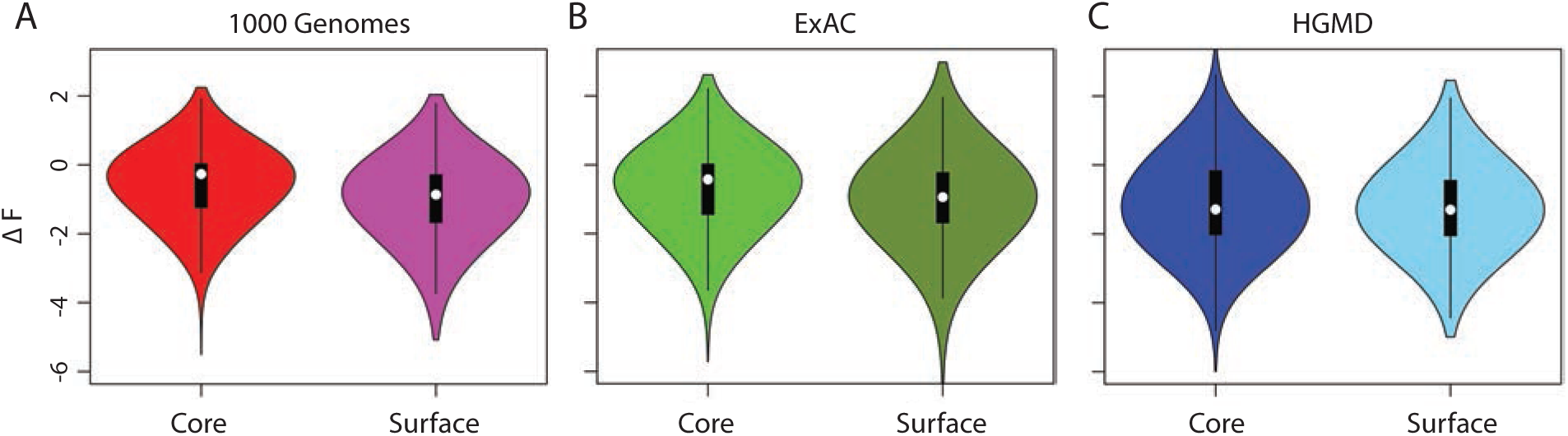
Differential effects of “benign” and disease-associated SNVs on the localized frustration of minimally frustrated residues in the non-mutated (i.e., native) state. Violin plots showing ΔF distributions associated with SNVs affecting the core or surface, with SNVs taken from *A)* 1000 Genomes, *B)* ExAC and *C)* HGMD. Comparison between ΔF distributions for core and surface residues of the 1000 Genomes and ExAC datasets indicate that favorable interactions of surface residues in the native states are highly disrupted upon mutation compared to core residues. Furthermore, ΔF in HGMD core residues were highly negative compared to 1KG and ExAC variants impacting core residues.

### Differential effects of rare and common SNVs on localized frustration

In population-level studies, SNVs with lower minor allele frequencies (MAF) are generally interpreted as being more likely to be deleterious than SNVs with higher MAF values. Thus, within the set of benign SNVs provided in the 1000 Genomes and ExAC SNVs, MAF may be used as an approximation for varying degrees of selective constraint. This prompted us to compare the rare and common SNVs induced ΔF distribution for minimally frustrated core and surface residues. Consistent with our earlier observations regarding benign SNVs, we found larger disruptions to favorable local interactions in surface residues relative to core residues (Figure 3A). However, this disparity was slightly more pronounced for rare SNVs compared to common SNVs. This observation was consistent for the 1000 Genome (Figure 3A) and ExAC datasets (Figure 3B) (*with p-value < 2e-16 from two-sample Wilcoxon test*). Furthermore, using both of these datasets, we observed that greater ΔF associated with the introduction of SNVs (in either the positive or negative directions) tend to be associated with lower MAF values (Figures 3C, top & bottom panels). This trend is observed for SNVs that occur on both the surface and within the core.

**Figure 3:**
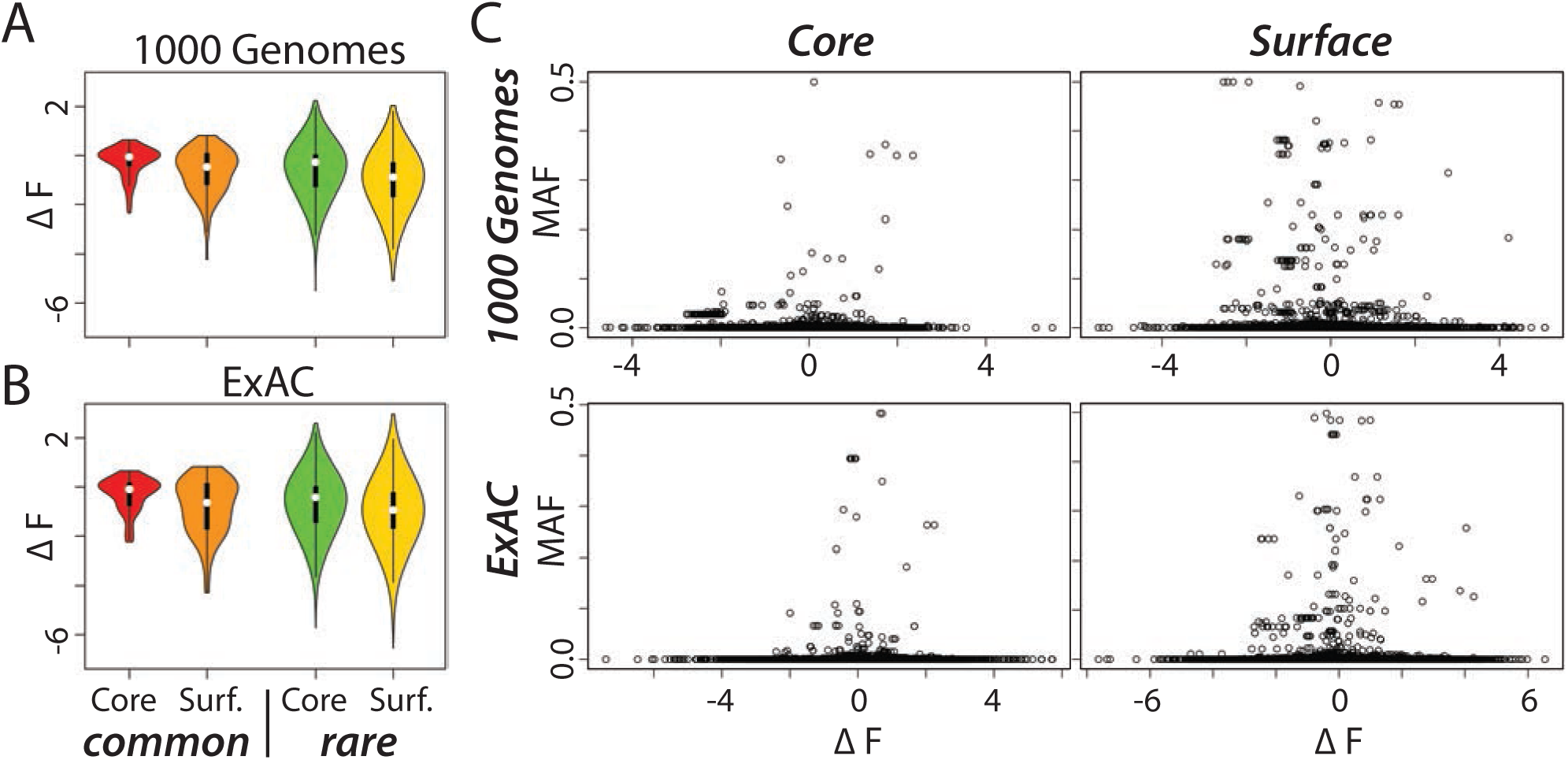
Common and rare SNVs differentially influence ΔF profiles of minimally frustrated core and surface residues. Violin plots show ΔF distributions induced by common and rare variants present in the 1000 Genomes *(A)* and ExAC *(B)* datasets (shown are the effects on minimally frustrated core and surface residues). Rare variants in both datasets lead to more substantially negative ΔF values compared to common variants. *C)* Scatter plots of ΔF values indicate that more extreme ΔF values (in either the positive or negative direction) tend to be associated with lower-MAF (i.e., rare SNVs).

### Differential effects of benign and disease-associated SNVs in different evolutionary contexts

We also examined the local perturbations induced by disease-associated and benign SNVs originating in conserved and variable regions of the genome. We plotted distributions for the ΔF values for the surface and core residues (Figure 4). We observed that benign SNVs originating in both conserved and variable regions of the genome had similar effects on minimally frustrated core residues (Figure 4A & 4B). This observation was true for the surface residues as well. In contrast, disease-associated SNVs intersecting with conserved and variable genomic regions lead to variable ΔF values for surface residues (*p-value = 0.00031 from two-sample Wilcoxon test*). This disparity is even more pronounced in core residues (*p-value = 3.298e-08 from two-sample Wilcoxon test*) (Figure 4C).

**Figure 4:**
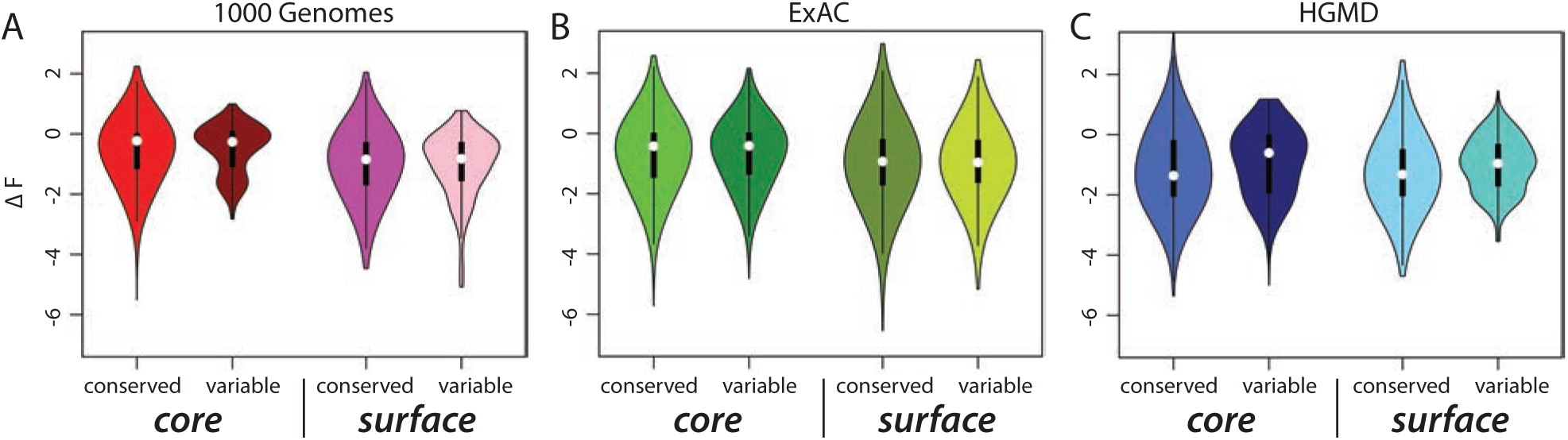
Comparisons between ΔF distributions associated with “benign” and disease-associated variants on evolutionarily conserved and variable residues. Violin plots depicting ΔF distributions introduced by *A)* 1000 Genomes, *B)* ExAC and *C)* HGMD variants, respectively. ΔF distributions associated with HGMD SNVs indicate larger disruption of conserved core residues compared to variable residues. In contrast, for the 1000 Genomes and ExAC datasets, no significant difference in ΔF distributions was observed for conserved and variable core residues (the same was true for surface residues).

### Differential effects of SNVs on driver and passenger genes

One of the most important challenges confronting the cancer genomics community involves discriminating between highly deleterious driver SNVs and the large number of neutral passenger SNVs that naturally arise over the course of tumor progression ^46^. As part of these efforts, a large number of cancer actionable genes have been curated in recent years. We applied our framework to evaluate the effects that somatic cancer SNVs have on driver genes ^47^, cancer-associated genes (CAGs) ^48^, and non-cancer associated genes (non-CAGs) in the context of frustration. We mapped the somatic pan-cancer SNVs that intersect these three distinct gene categories onto protein structures. We then evaluated the ΔF distributions in all three categories.

As with benign SNVs, we observed that somatic SNVs impacting CAGs and non-CAGs lead to greater disruptions in minimally frustrated surface residues relative to core residues (*p-value < 2.2e-16 from two-sample Wilcoxon test*) (Figure 5). Moreover, this variability in ΔF distributions between core and surface residues was more pronounced among non-CAGs compared to CAGs (Figure 5). In contrast, SNVs that impact driver genes lead to larger disruptions in favorable localized interactions for surface and core residues (*p-value < 2.2e-16 from two-sample Wilcoxon test*) (Figure 5) compared to CAG core and surface residues.

**Figure 5:**
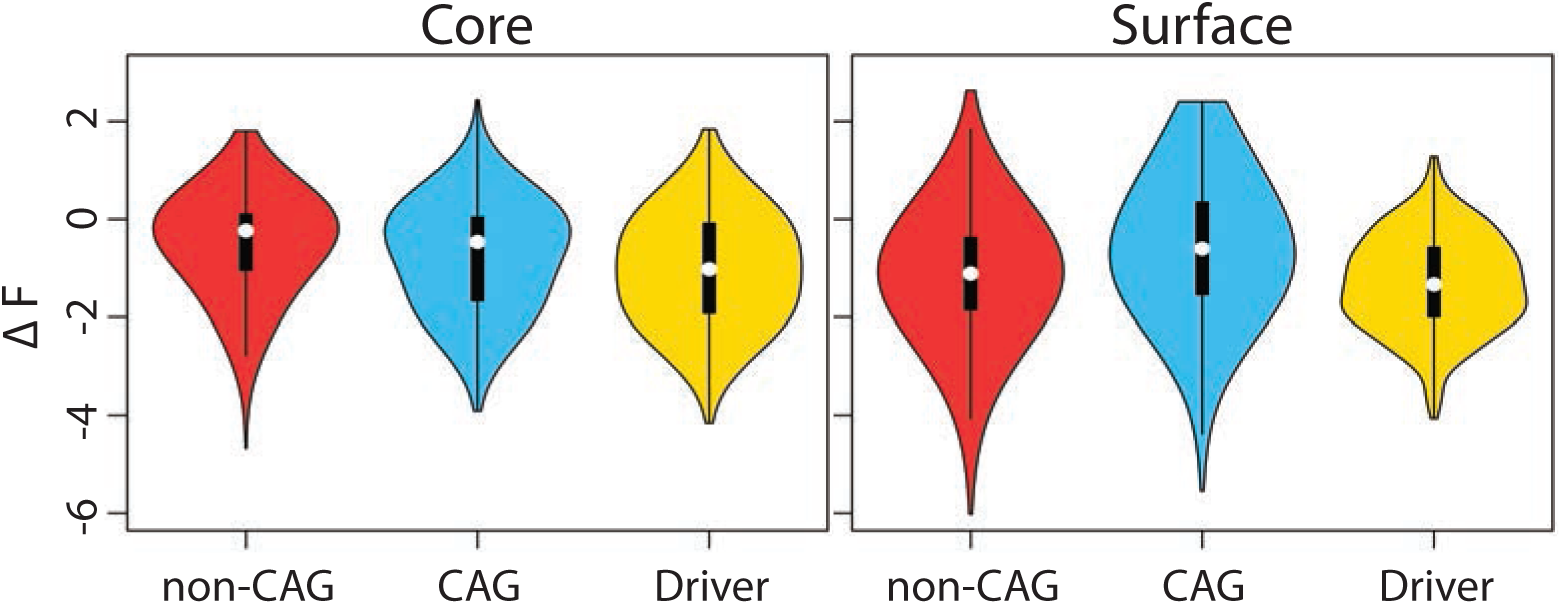
Comparisons between ΔF distributions associated with driver and passenger genes. *Left)* Violin plots showing ΔF distributions associated with somatic SNVs affecting *non-cancer associated genes (non-CAG), cancer associated genes (CAG) and driver genes* encoding core and surface residues. Somatic SNVs affecting core residues of driver genes lead to a more substantially negative ΔF values compared to those in CAG and non-CAG proteins. *Right)* On the contrary, SNVs in CAGs and non-CAGs disrupt favorable interactions of the surface residues to a larger extent compared to their core residues.

### Differential effects of SNVs on oncogenes and tumor-suppressor genes

Cancer driver genes are classified as oncogenes and tumor suppressor genes based on their mutational pattern and their mode of inducing tumorigenesis ^47^. Oncogenes are marked by recurrent SNVs within the same gene loci across different cancer types, and are believed to drive cancer progression through gain-of-function (GOF) mechanisms. In contrast, a tumor suppressor gene generally contains protein-truncating mutations or SNVs that are scattered throughout the gene, and they are believed to facilitate cancer progression through loss-of-function (LOF) mechanisms. This line of thinking is guided by the idea that LOF variants often act by destabilizing the protein (Figure 6C, left panel), whereas GOF variants may impact protein-protein interaction interfaces (by reducing specificity for binding partners) or negatively affect auto-regulatory sites on the protein, many of which are on the surface (Figure 6C, right panel).

**Figure 6:**
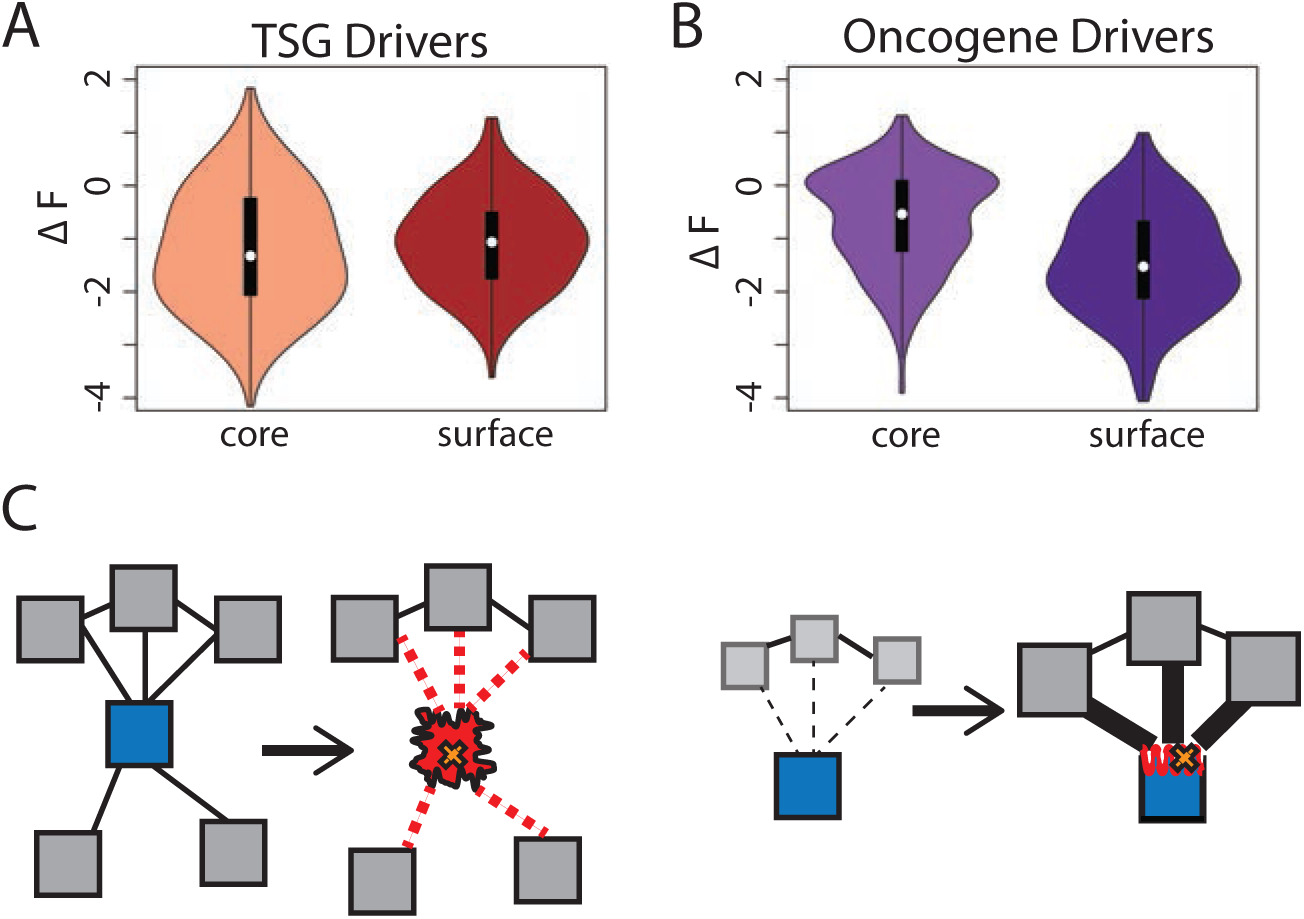
Differential impacts on ΔF distributions associated with of SNVs on driver and passenger genes. Violin plots demonstrating ΔF distributions associated with SNVs tumor suppressors *(A)* and oncogenes *(B)*. SNVs in tumor suppressors (TSGs) lead to larger disruptions for minimally frustrated core compared to surface residues. However, SNVs affecting oncogenes are associated with larger ΔF values for the surface compared to core residues. *C)* These observations suggest a potential model in which SNVs in TSGs act by disrupting the hydrophobic core of a protein and drive cancer progression through LOF mechanisms *(left)*. In contrast, SNVs in oncogenes may facilitate non-specific binding by changing surface residues and drive cancer through GOF events *(right)*.

In order to evaluate the extent to which such effects manifest in our set of tumor-suppressor genes and oncogenes, we applied the frustration framework to evaluate changes in local perturbation when SNVs impact these distinct categories of driver genes (Figure 6A& 6B). We observed that SNVs affecting TSGs induce stronger perturbations in minimally frustrated core residues relative to surface residues (*p-value = 8.15e-2 from two-sample Wilcoxon test*) (Figure 6A). In contrast, SNV affecting oncogenes induces greater ΔF values within minimally frustrated residues in the surface relative to core residues (*p-value = 2.2e-16 from two-sample Wilcoxon test*) (Figure 6B). Moreover, SNVs impacting *oncogenes* lead to larger disruptions in favorable local interactions compared to *TSGs* for minimally frustrated surface residues (*p-value = 5.0e-4 from two-sample Wilcoxon test)*. However, SNVs impacting TSGs lead to greater disruption in favorable local interactions compared to oncogenes affecting driver SNVs in core residues (*p-value = 6.306e-15 from two-sample Wilcoxon test*).

## Discussion

In the last decade, tremendous improvements in sequencing and structural biology techniques have lead to growth in genomic variation and three-dimensional structural data for various proteins. This concomitant growth in the sequence and structural space provide us with an ideal platform to investigate the impact of genomic variants on protein structure. The objective of these studies is to gain mechanistic insights into the origin of various diseases, as well as design effective drug targets for them. Prior studies in this direction were limited due to lack of genomic variation and structural data. Moreover, these studies primarily focused on investigating the impact of SNVs on the *global* stability of protein structure. However, many experimental studies have clearly indicated causal role of SNVs induced local perturbation in various diseases. In this work, we repurpose the concept of localized frustration, originally introduced in protein folding studies to quantify SNV-induced local perturbations. The frustration index of a residue quantifies the presence of favorable/dis-favorable local interactions in the protein structure compared to a random molten globule structure.

In this study, we employed an extensive catalogue of benign (∼5.7 million) and disease-associated (∼0.76 million) SNVs. The benign SNV dataset comprised of SNVs from the 1000 Genome project (phase 3) and the ExAC project. In contrast, HGMD SNVs and pan-cancer somatic SNVs constituted our disease-associated SNV dataset. We mapped ∼0.2 million benign and disease-associated SNVs onto ∼10K high-resolution protein structures. Subsequently, we compared the impact of benign and disease SNVs on the frustration profile of minimally frustrated residues in various protein structures. The ΔF distributions indicated that both benign and disease SNVs disrupt minimally frustrated surface residues to similar extents. However, the mechanistic difference between benign and disease SNVs can be attributed to their impact on the local environment of core residues. Within the core, disease-associated SNVs result in more severe perturbations to local interactions relative to those introduced by benign SNVs. These local disruptions are propagated throughout the core and, in turn, drive the deleteriousness of various disease-associated SNVs.

Furthermore, we quantified the influence of rare and common SNVs present in healthy human population on the frustration profile of affected protein residues. We observed that rare SNVs lead to larger local perturbation of minimally frustrated surface residues compared to common SNVs. This observation is intuitively consistent as one would expect rare SNVs to have grater impact on protein stability. In addition, we also investigated the differential impact of SNVs intersecting conserved regions compared to variable regions of the genome. The distinction between conserved and variable regions of the genome was based on GERP scores, which quantifies a cross-species conservation score on each nucleotide position of the genome. This cross-species conservation analysis indicated that there is no disparity between ΔF associated with benign SNVs fixated in conserved and variable regions. This lack of disparity can be attributed to the absence of significant local perturbations induced by benign SNVs, which do not compromise the overall stability of protein structure. In contrast, for disease SNVs originating in conserved and variable regions of the genome, we observe significant differences in ΔF values. This is consistent with prior studies, which indicate that the deleteriousness of an SNV is more pronounced when SNVs impact functionally important conserved regions of the genome compared to variable regions of the genome.

In addition to studying disease variant in general, tremendous progress in next generation sequencing has lead to unprecedented efforts to characterize cancer genome. Large efforts have been invested in discriminating between driver and passenger SNVs. Driver SNVs are known to play important roles in driving cancer progression. Motivated by this, we examined the influence of SNVs emanating in driver and passenger genes. Specifically, we studied these effects in the context of the local stability of protein structure. Our analysis indicated that SNVs influencing non-actionable genes (non-CAGs) and indirectly actionable genes (CAGs) lead to greater perturbations of surface residues compared to core residues. In contrast, SNVs that impact driver genes have similar affects on ΔF values in core and surface residues. These observations further reiterate our earlier conclusion that the deleteriousness of a given SNV is determined by its ability to perturb the local interactions of core residues. These local perturbations further propagate through the core to completely destabilize the protein structure.

Furthermore, cancer driver genes are often classified as oncogenes and tumor suppressor genes based on their mode of cancer progression. SNVs in oncogenes lead to cancer progression through GOF mechanism, whereas SNVs impacting tumor suppressor genes contribute to cancer growth through LOF events. These two distinct mode prompted us to closely inspect SNVs originating in oncogenes and TSGs. We compared the ΔF profile for residues influenced by these two distinct categories of SNVs. we observed that SNVs in oncogenes and TSGs generate greater ΔF values for surface and core, respectively.

Comprehensive catalogues of genomic variations from large-scale genomics project have clearly established the important role of disease-associated and rare variants in human populations. We foresee further growth in genomic variation datasets as large-scale genomic consortium projects such as International Cancer Genomics Consortium (ICGC), The Pan-Cancer Genome Atlas (PANCAN Atlas), UK10K project and Mendelian genomic program will continue to decipher mutational landscape of human genomes and exomes. Similarly, advancement in electron microscopy, NMR, small angle X-ray scattering and other biophysical techniques will further increase the availability of protein structural data. These expanding knowledge bases of genomic variation and structural biology will facilitate integrative studies to gain mechanistic insight in disease progression and design effective drugs for disease treatment. In this work, we demonstrate the role of localized frustration as a metric to quantify and investigate the influence of genomic variants on protein structures. The proposed framework is a logical extension to some of the earlier studies, which primarily employed global metrics such as folding free energy changes to quantify the affects of genomic variants. We strongly believe that combination of these global and local metrics, along with sequence features, will help us elucidate the mechanism as well as predict the impact of genomic variations in disease and and healthy human populations.

## Methods

### SNV Datasets

We utilized a comprehensive catalogue of SNVs from various resources. Our SNV dataset is divided into two broad categories (benign and disease-associated) (S1A). The benign set comprises of SNVs reported in The 1000 Genome Project (phase 3) ^7^ and The Exome Aggregation Consortium ^10^. Disease-associated dataset included SNVs from the Human Genome Mutational Database (HGMD) ^5^ and pan-cancer dataset ^49^ comprising of publicly available somatic SNVs from The Cancer Genome Atlas (TCGA) ^50^, The Catalogue of Somatic Mutations in Cancer (COSMIC) ^51^ and the SNV dataset available from Alexandrov *et. al* ^52^. SNVs from the pan-cancer dataset were further sub-classified (driver and passenger sets) based on whether they are mutating a driver or passenger gene. Driver genes were curated from the Vogelstein et. al. ^47^, where they distinguish between driver and passenger genes based on mutational patterns. They define a driver gene as an oncogene if the SNV is recurrent at the same gene loci, whereas tumor suppressor genes (TSG) are mutated throughout their length. Similarly, we sub-classified passenger genes into cancer-associated genes (CAGs) and non-cancer associated genes (non-CAGs). CAGs included genes from the cancer gene census (CGC)^53^ and a curated list of 4050 genes from a previous study ^48^. Furthermore, we removed any driver gene present in the CAG dataset. The remaining set of genes impacted by pan-cancer SNVs constituted our non-CAG dataset.

### Workflow to calculate frustration

As mentioned earlier, we investigated the impact of different categories of SNVs on the local stability of various protein structures. We utilize the ΔF values of mutated residues to quantify SNVs induced local perturbation. Quantifying ΔF involves three steps: a) mapping SNVs onto the affected three-dimensional structure, b) generating the homology model of the mutated structure, and c) evaluating the ΔF of mutated residue in the native and mutated conformations.

To map SNVs onto protein structures, the Variant Annotation Tool (VAT)^54^ was applied to annotate our curated catalogue of SNVs. This annotation includes the gene and transcript names, residue position in the protein sequence, as well as the original and mutated residue identity. We then integrated VAT annotation with the biomart^55^ derived human gene and transcript IDs to map the SNV on to specific PDB structures. We restricted this SNV mapping scheme to high-quality structures with resolution values that were better than 2.0 Angstrom. Following the SNV mapping to PDB structures, we generated models of the resultant mutated structures by applying homology modeling using the mutated protein sequence and native protein structure as input to modeler ^56,57^.

Finally, we quantify the frustration index of the mapped residue in the native structure as well as in the mutated model of the protein. Briefly, the residue level localized frustration index ^45^ quantifies the degree to which that amino acid favorably contributes to the energy of the system relative to all 20 possible amino acids at that position:
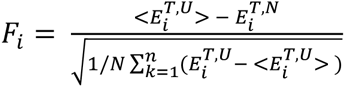, where 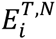 is the total energy of the protein in the native state. The total native energy is calculated using a function that includes an explicit water interaction term,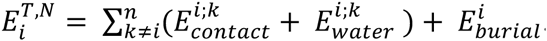 This function, termed the associated water-mediated (AWM) potential [44], describes the energies associated with direct interactions between residues *i* and *k* (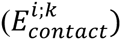 as well as those with water-mediated interactions between residues *i* and *k* 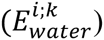 and energy term 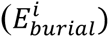 associated with the burial of the residue. The average energy of the decoy conformations (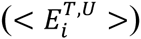) is generated by mutating the original residue *i* to each of the alternative possible nineteen residues. The AMW potential includes different parameter values for different residues, so the decoy energies calculated vary based on the identity of the mutated residue.

In Figure 1, we demonstrate an example case in which replacing isoleucine at a particular locus within ubiquitin (PDB ID 1UBQ) with a tyrosine. Shown on the left (in green) is the native (i.e., wild-type structure). The vertical axis designates the different energies that would result when the residue at this locus is mutated to each one of the other 19 amino acids. Specifically, these 19 decoy energies are only calculated by changing the parameter values that are specific to each amino acid within the potential function (note that the structure is not altered or minimized in any way). In the sense that these energies are calculated in the context of the structure that is otherwise identical to the wild-type X-ray structure, the energy distribution shown at left represents the energies in “native structure”. The dotted line represents the mean value among all of the 20 energy values associated with the various amino acids. In this case, the energy computed using the wild-type residue (ILE) is substantially lower than this mean value (rendering ΔE_nat_ greater than 0). Because ΔEn_at_ is greater than 0, this wild-type isoleucine is said to be “minimally frustrated”.

This same protein is known to contain a disease-associated SNV at locus 31. Specifically, the disease-associated change occurs when the isoleucine is mutated to tyrosine. To quantify ΔF in this example, we first introduce the tyrosine at locus 31 *in silico*, and then use Modeller to generate a model of the mutated structure (shown at right, in orange). Thus, we now not only change the type of residue at locus 31, but also the configuration of the entire protein; the structure is that said to be “non-native” (the relative energy values given on the horizontal axis may thus become redistributed slightly). In this new energy landscape, the energy associated with the residue at the mutated locus 31 is higher than the mean energy among all 20 amino acids within the modeled structure (ΔE_mut_ < 0), suggesting that the mutated residue is “maximally frustrated”. We are primarily interested in the ΔF between these two states. This value is proportional to the difference between ΔE_mut_ and ΔE_nat_. (ΔE_mut_ - ΔE_nat_ = ΔΔE) Here, ΔΔE is less than 0, suggesting that the frustration is higher in the mutated structure than that of the wild type.

## Acknowledgments

We acknowledge support from the NIH and from the AL Williams Professorship funds. We thank Diego Ferreiro for helpful discussion and sharing the original source code for localized frustration calculations. We also acknowledge help of Anurag Sethi and Suganthi Balasubramanian for providing valuable feedbacks for improving the manuscript. The authors would like to thank the Exome Aggregation Consortium and the groups that provided exome variant data for comparison. A full list of contributing groups can be found at exac.broadinstitute.org/about

